# North American bird occupancy dynamics attributed to climate and land use change

**DOI:** 10.64898/2026.08.24.746675

**Authors:** Katrin Schifferle, Natalie J. Briscoe, Guillermo Fandos, Stefanie Heinicke, Christopher P. O. Reyer, Inga J. Sauer, Mark C. Urban, Damaris Zurell

## Abstract

Evidence is accumulating that global change is altering species distributions. Yet, detailed knowledge is missing about the relative and joint contribution of different drivers to observed species responses.

Here, we implemented an impact attribution framework based on counterfactual simulations to assess the impact of climate and land use change on occupancy dynamics of North American breeding birds. We used a Bayesian framework to fit process-explicit dynamic occupancy models to long-term survey data for 159 species from 1995 to 2019, and quantified predictive performance using spatial and temporal cross-validation. We then assessed the relative importance and effect direction of climate and land use change while accounting for model predictive accuracy.

Results indicate that climate change negatively affected 90 % of the species and land use change negatively impacted 96 %. Climate change emerged as more important than land use change for driving changes in occupancy across species. Remarkably, the effects of both drivers were mostly antagonistic rather than acting additively or synergistically. Climate was the most important driver for bird communities in the western USA, while land use change dominated in the southeast, and combined climate and land use change in the northeast.

Our analysis demonstrates that recent changes in North American bird distributions are shaped by multiple global change drivers acting in concert. The effect of recent climate and land use change were mostly antagonistic, and thus trends in bird occupancy dynamics could not be understood by studying the impact of those drivers in isolation. By disentangling the effects of climate and land use change on biodiversity trends, impact attribution approaches can improve our understanding of global change impacts and can support conservation planning and more accurate and realistic projections of biodiversity response to global change.

## Introduction

Human changes to the Earth system have increased rapidly, and these disturbances have negatively affected biodiversity worldwide (Steffen et al., 2015; IPBES, 2019). Currently, the most important direct global driver of biodiversity change is land and sea use change, but climate change impacts are accelerating (IPBES, 2019). Drivers often co-occur (Bowler et al., 2020), and hence they could interact in their impacts on biodiversity (Sala et al., 2000). Whether species benefit from environmental change or are disadvantaged depends not only on their level of exposure to change, but also on species-specific sensitivity and adaptive capacity (Dawson et al., 2011; Williams et al., 2008). Although evidence is accumulating that the range limits of many species are expanding poleward and uphill (Parmesan & Yohe, 2003; Martins et al., 2024) in line with the assumption that global warming is a major driver, many species demonstrate idiosyncratic or limited range changes over the last decades (Hockey et al., 2011; Lenoir & Svenning, 2015; Currie & Venne, 2017; Martins et al., 2024; Zurell et al., 2024). These divergent patterns underscore the need to identify the winners and losers of global change and to link these outcomes to specific drivers to generate more accurate and realistic projections of biodiversity change and to implement efficient measures to protect biodiversity (Bowler et al., 2020; Clement et al., 2019; Schrodt et al., 2025; Sirami et al., 2017).

Assessing the relative contributions of different drivers to observed changes in a system is termed attribution (Ara Begum et al., 2022). Recently, scientists have argued for the critical need for methods to attribute changes in ecological systems to human-induced drivers (Gonzalez et al., 2023; Dudney et al., 2025; Schrodt et al., 2025). Climate impact attribution assesses the impacts of climate change on coupled natural-human systems (Frieler et al., 2024) and, according to the Working Group II definition of the Intergovernmental Panel on Climate Change (IPCC), can be achieved by quantifying the difference between the observed state of a system and a counterfactual baseline, i.e. the state of a system in the absence of climate change (O’Neill et al., 2022). To date, few studies have attributed the drivers of change in ecological systems in line with this definition. Examples include the attribution of bird abundance change to climate change impacts and human pressures (Kotz et al., 2025) as well as the attribution of range shifts and expansions of infectious diseases affecting humans in Europe (Erazo et al., 2024) and trees in the Sierra Nevada (Dudney et al., 2021) to climate change. While these attribution studies employ static statistical models, models that explicitly account for temporal dynamics and non-equilibrium responses may provide more reliable estimates of species responses under ongoing environmental change (Briscoe et al., 2021; Zurell et al., 2016), which is particularly relevant for attributing impacts of global change drivers.

Dynamic occupancy models (DOMs) match these criteria. They explicitly represent changes in species distributions through colonisation and extinction processes, account for imperfect detection, and relax the equilibrium assumption (MacKenzie et al., 2003). DOMs estimate initial occupancy and subsequent colonisation and extinction rates from species detection/non-detection data, with these processes potentially modelled as a function of environmental covariates. DOMs have been demonstrated to capture changes in temporal population dynamics particularly well (Briscoe et al., 2021) and their intermediate level of complexity makes them well suited to model occupancy dynamics for many species and across large areas (Kelleher et al., 2025). Moreover, DOMs provide output that can be rigorously validated based on species observations, which strengthens confidence in counterfactual simulations as a basis for impact attribution analyses.

Here, we combined DOMs with an impact attribution framework to assess the relative importance of climate and land use change for observed occupancy dynamics of breeding birds in the conterminous USA. The North American Breeding Bird Survey (BBS) provides a rich dataset with annual surveys. It started in 1966 in the eastern USA and has expanded in spatial coverage in more recent decades (Ziolkowski et al., 2024). Evidence is accumulating that North American breeding bird populations are undergoing major changes. Across species, multiple independent monitoring methods have revealed substantial declines in abundance over the past decades (Rosenberg et al., 2019) and taxonomic diversity since the 2000s (Jarzyna & Jetz, 2017). While land use change is considered a main driver of changes in bird communities (Barbet-Massin et al., 2012; Lemoine et al., 2007), peak abundances have shifted northwards, particularly in western North America, which suggests climate change might also be a likely driver of change (Martins et al., 2024).

In this study, we therefore focused on climate and land use change as potential drivers of past bird occupancy change. We used single-species DOMs combined with counterfactual climate and land-use scenarios to attribute observed occupancy dynamics between 1995 and 2019 to climate change, land use change, and their interaction. Based on this analysis, we assessed 1) the proportion of climate and land use change winners and losers across North American breeding bird species and 2) the relative importance of climate change, land use change and the combination of both drivers for the observed occupancy dynamics.

## Methods

### Overview

We modelled occupancy over time with dynamic occupancy models (DOMs; MacKenzie et al., 2003). In these models, occupancy change is derived from estimates of initial occupancy probability and subsequent local colonisation and extinction probabilities. We fitted single-species DOMs based on annual bird observations and annual data of observed climate and reconstructed land use spanning the conterminous USA over the 25-year period 1995-2019. Model predictive performance was assessed using spatial and temporal cross-validation and only models with acceptable predictive performance in both dimensions were carried forward to the attribution stage. We used these models to simulate occupancy time series under observed climate and reconstructed land use change (“factual simulations”; Figure 1a), as well as under counterfactual scenarios of no-climate change, no-land use change and neither climate nor land use change (“counterfactual simulations”; Figure 1b). To identify winners and losers of change, we first compared linear temporal trends in occupancy across scenarios. Second, we quantified the relative importance of climate and land use change in driving variation in species occupancy over time. To do this we calculated the difference in prediction error of factual and counterfactual simulations, which controls for differences in predictive performance between models (Figure 1c).

**Figure 1.**
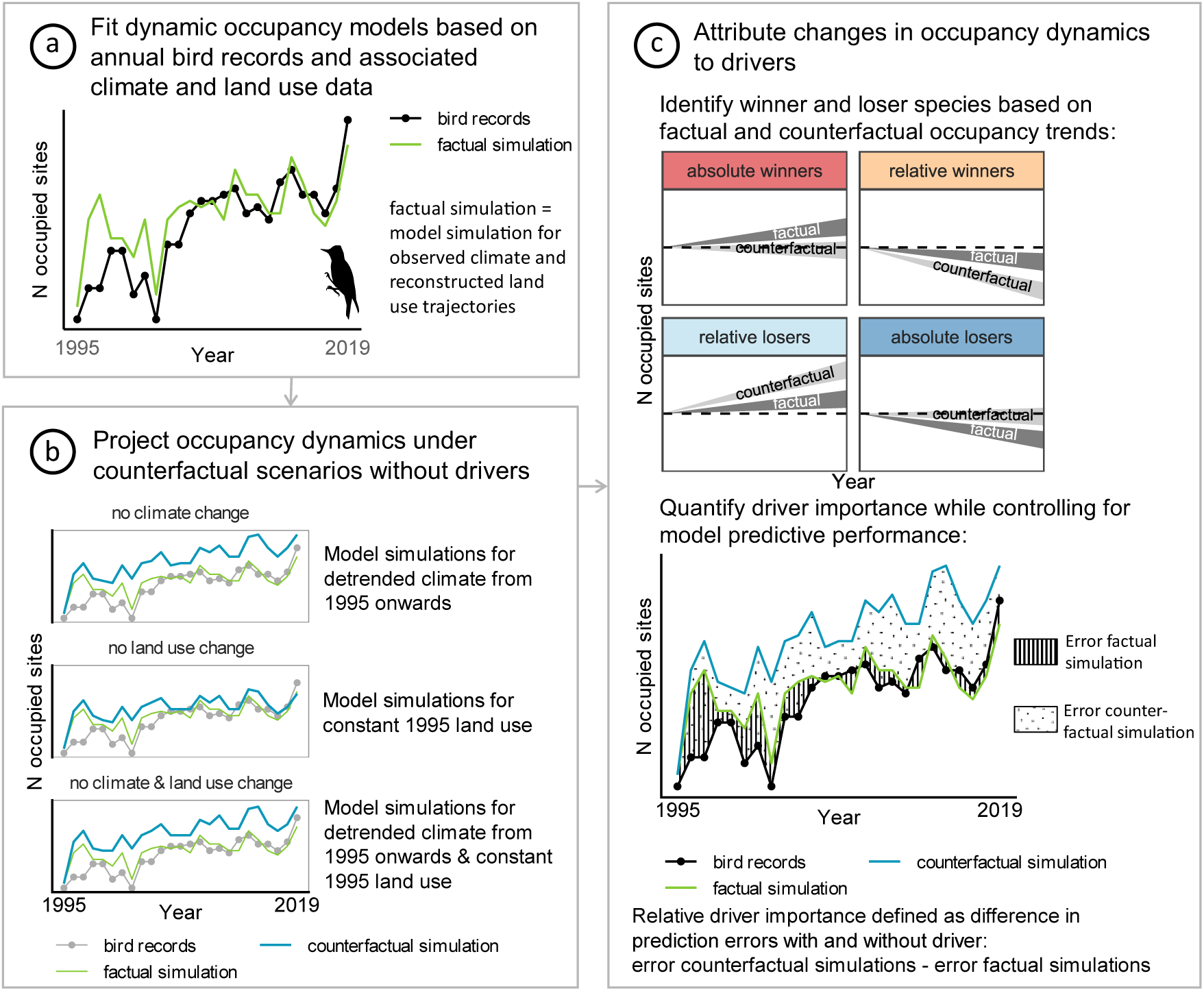
Attribution framework used to disentangle the importance of climate and land use change for changes in occupancy dynamics of 80 North American breeding bird species from 1995 to 2019. a) We first fitted models of occupancy dynamics based on bird observations from the North American Breeding Bird Survey and observed climate and reconstructed land use. With these models we simulated occupancy dynamics under climate and land use change (“factual simulations”). b) We used the models with acceptable predictive performance to then simulate occupancy dynamics for scenarios without climate change, without land use change and with neither climate nor land use change (“counterfactual simulations”). c) To attribute the effects of climate and land use change we assessed the difference between model simulations with and without the respective driver(s). We first compared the occupancy trend in factual and counterfactual simulations to identify winner and loser species of climate and land use change (upper panel in c adapted from Langhammer, et al., 2024). We then quantified the importance of each driver as the difference in predictive accuracy between factual and counterfactual simulations (lower panel in c). Bird records and simulations are shown for yellow-bellied sapsucker (*Sphyrapicus varius*). Bird silhouette from PhyloPic (https://www.phylopic.org/; T. Michael Keesey, 2023), silhouette was made by Andy Wilson (CC0 1.0).

We used R (R Core Team, 2025) for all analyses, except for creating counterfactual climate data (see below). The R code and the complete list of R packages used to reproduce this work can be found in the supplemental material (https://doi.org/10.5281/zenodo.22080744).

### Data processing

#### Bird observations

We used data from the North American Breeding Bird Survey (BBS, Ziolkowski et al., 2024). The BBS contains annual bird counts along randomly established transects (routes). Each route follows secondary roads and is roughly 40 km long. Approximately every 800 m observers stop along the road and conduct a three-minute point count of all birds seen and heard within a 400 m radius. In total, observers record observations at 50 stops per route. The surveys are performed once per year at the height of the breeding season, mostly in June, by skilled citizen scientists and begin 30 minutes before sunrise. We used the dataset in which counts are summarized at 10-stop intervals since counts at single stops were only reported from 1997 onward. We imported the data into R using functions of the bbsAssistant R package (Burnett et al., 2019) and converted count data into detection/non-detection data required by DOMs.

##### Route selection

Our aim was to compile a dataset that contains information across the entire breeding ranges of many species, representing range edges as well as range interiors, and that covers as long of a period as possible with consecutively surveyed years to capture occupancy dynamics across the range. Selection criteria for routes included in this study are listed in Table S1. We focused on the time period 1995 to 2019 because a large number of routes had been surveyed nearly annually during this period, and because our chosen annual climate and land use datasets were not available for years after 2019. We selected only routes that had been surveyed in at least 20 of the 25 years (i.e. 80 %). To reduce bias in the spatial coverage of the USA, we thinned the routes such that the minimum distance between two route centroids was 100 km (twice the spatial resolution of the environmental data, see below) and only kept a maximum of 30 routes per Bird Conservation Region (Jarzyna & Jetz, 2017). The final selection contained 539 routes (Figure S 1).

##### Species selection

We summarised records of subspecies differentiated in the BBS at the species level. We then removed all records of birds that had not been identified at the species level as well as records of hybrids. We excluded all nocturnal and water-related species since these are not well sampled by the BBS methods (Harris et al., 2018). We further excluded very rare or widespread species for which obtaining informative models is challenging due to a small number of colonisation and local extinction events in the data. We excluded rare species that were detected on fewer than 50 different routes over the study period, as well as very widespread species with fewer than 50 routes on which they were not detected. The resulting dataset included 192 common breeding bird species.

#### Environmental data

Climate and land use data were downloaded from the Inter-Sectoral Impact Model Intercomparison Project (ISIMIP) Repository (https://data.isimip.org/, last access: 30-01-2026) for ISIMIP phase 3a. ISIMIP3a offers a framework to consistently attribute the observed impacts of climate change across different sectors (Frieler et al., 2024). Climate and land use data have 0.5° spatial resolution which corresponds to a cell size of approximately 45 km in the conterminous USA, projected with Albers equal area projection. This roughly corresponds to the BBS route length of 40 km, which is the unit of our analysis. We matched routes to environmental data based on the locations of the routes’ centroids.

##### Climate

We used daily GSWP3-W5E5 climate data of ISIMIP3a (Lange et al., 2025). For our study period from 1995-2019, these are based on ERA5, which is a state-of-the-art reanalysis product that provides global datasets of climate variables by combining observations with modelling techniques (Hersbach et al., 2020). To fit the DOMs, we used monthly means of daily minimum and maximum temperature and precipitation (‘obsclim’ scenario), from which we calculated annual bioclimatic variables using the R package dismo (Hijmans et al., 2023). We defined the annual period from June of the previous year to May of the current year to capture the climate that preceded the current breeding season, including any beneficial or detrimental effects of autumn and winter of the previous year (Barbet-Massin & Jetz, 2014). Since we found discontinuities in variables combining temperature and precipitation information (bio8, bio9, bio18 and bio19) in the conterminous USA, we discarded them as recommended by Booth (2022). Since the quarterly variables of the bioclimatic variables, especially those referring to the driest or wettest quarter of the year, can refer to different seasons in different regions of the USA, they may affect birds during different periods of their annual cycle. Therefore, we additionally calculated mean temperature and precipitation values of each season (spring: March to May, summer: June to August, autumn: September to November, winter: December to February) as quarterly variables that have a fixed temporal relationship with the birds’ annual cycle.

##### Land use

We used ISIMIP3a’s land use input dataset that contains for each year and cell the fraction covered by each land use class (Volkholz & Ostberg, 2024). It is derived by interpolation from the Land-Use Harmonization (LUH2) dataset (Hurtt et al., 2020). For model fitting we considered the categories “forest and natural vegetation”, “managed pastures and rangeland”, “urban areas” and “five crop types”. From these categories we initially considered the following land use classes: primary non-forests, secondary forest, secondary non-forests, managed pastures, rangeland, urban areas and annual crops. As annual crops, we summarised C3, C4 and C3 nitrogen fixing crops (Naimi et al., 2022). We did not consider perennial crops and primary forests since relevant fractions occur only in small parts of the conterminous USA.

##### Counterfactual environmental data

Since we were interested in the impact of climate change that had accumulated over the 25 year study period, we generated counterfactual climate data that are detrended from 1995 onwards rather than using the counterfactual data available from ISIMIP, in which the effect of climate change since 1901 was removed (Frieler et al., 2024). We used the ATTRICI (ATTRIbuting Climate Impacts) command line tool (https://github.com/ISI-MIP/attrici). The ATTRICI approach is tailored to attributing impacts of climate change, irrespective of its cause (Mengel et al., 2021). ATTRICI constructs counterfactual climate data without long-term trends from factual climate data while preserving internal data variability. Based on the resulting counterfactual climate data, we calculated bioclimatic variables and seasonal averages in the same way as for the factual data.

For land use, we followed ISIMIP’s rationale and fixed land use at a specific year. In our case, 1995 (Volkholz & Ostberg, 2024), to maintain a constant land use at a level that would provide a counterfactual comparison. This way, we quantify the impact of changes in direct human forcing by comparing simulations for factual human forcing to simulations for constant human forcing fixed at certain years (Frieler et al., 2024). We expected the direction of any bias resulting from considering interannual variability in counterfactual climate data, but not in land use data, to be small because land use fractions in the conterminous USA showed limited interannual variability over 1995–2019.

### Model fitting

### Dynamic occupancy models (DOMs)

To simulate the occupancy dynamics of breeding bird species under climate and land use scenarios, we fitted single-species DOMs (MacKenzie et al., 2003). These models estimate initial occupancy probability and subsequent colonisation and extinction probabilities. Based on these estimates, occupancy dynamics are simulated as a Markov process, i.e. the occupancy at one time step depends on the occupancy in the previous time step as well as on the respective colonisation and extinction probabilities. Since DOMs require data from repeated surveys to estimate detection probabilities, we considered each route as a single location and five sections along a route, consisting of ten stops each, as replicate observations. By doing so, we defined occupancy at the route level, i.e. as the presence of a species along a BBS route during one breeding season (Royle & Kéry, 2007; Jarzyna & Jetz, 2017; Doser et al., 2023).

To assess the impacts of climate and land use change across species, we used the same set of covariates for each species. We selected variables that cover a substantial amount of environmental variation across the conterminous USA and that are not substantially correlated. This resulted in a selection of 15 variables, ten related to climate: annual mean temperature, mean diurnal temperature range, annual temperature range, temperature isothermality, precipitation seasonality, precipitation of driest month and mean spring, summer, autumn and winter precipitation; and five variables related to land use: primary non-forest, secondary non-forest, managed pastures, urban areas and annual crops. Details about the variable selection procedure and assumptions underlying the DOMs are listed in the Supplementary Material.

We estimated the process rates of colonisation and extinction based on annually resolved climate and land use data assuming that the climatic and land use conditions in the breeding area over the past twelve months affected local colonisation and extinction (Clement et al., 2016). As potential covariates for initial occupancy, we used the averages of the three years before our focal time period (1992 to 1994) to account for the cumulative impact of environmental conditions on site occupancy and to allow estimates of initial occupancy that are more robust with regard to annually varying environmental conditions. For all environmental covariates, we included linear and quadratic effects. To account for the fact that bird activity, and therefore likely also their detectability, varies with the time of the day (Clement et al., 2016), we modelled detection probability based on the route section (1-5) that we used as a proxy for time of day (surveys always start 30 minutes before local sunrise). All variables were standardized to having a mean of zero and a standard deviation of one.

We aimed to fit single-species models for each of the selected 192 species. For each species, we considered all routes within a buffer of 750 km around all presences recorded between 1995 and 2019 to fit the models. By restricting the considered non-detections, we expected to fit more meaningful models since not every part of the conterminous USA is accessible to every species regardless of climate and land use, due to dispersal constraints. A buffer size of 750 km is an arbitrary choice but seemed appropriate for the majority of species and to account for the Rocky Mountain as a major dispersal barrier, while considering that birds can have high movement ability. In a preliminary analysis, we tested a smaller buffer size of 250 km for a subset of species, but found no substantial differences in the predicted occupancy dynamics over time compared to a buffer size of 750 km. A 750 km buffer also allowed us to conduct a five-fold spatially blocked cross validation for model evaluation, since the resulting buffer area is large enough to be split into five blocks while maintaining a large enough number of detections and non-detections within each block to refit the models (see below).

We used a Bayesian approach to fit the models with the R package flocker (Socolar & Mills, 2023) which fits models in Stan via brms (Bürkner, 2017). We fitted the colonisation-extinction (“colex”) model and used logistic priors with location (mu) = 0 and scale (sigma) = 1 for the intercepts, which are uniform at the probability scale, and weakly informative priors with mean (mu) = 0 and standard deviation (sigma) = 2 for the coefficients (Northrup & Gerber, 2018; Socolar & Mills, 2023). We ran four chains with initially 1000 iterations of warmup followed by 1000 sampling iterations. We then checked diagnostics on the Markov Chain Monte Carlo (MCMC), using the R package bayesplot (Gabry & Mahr, 2024): we visually checked the trace plots and tried to refit the models with 2000 iterations of warmup followed by 2000 sampling iterations if either R-hat was > 1.02 for at least one parameter, or if the effective sample size or the effective sample size in bulk or tail was below 10 % of the total sample size or if there were divergent transitions. We encountered one or more of these issues for 14 of the 192 species. For nine species the issues remained after running more iterations. These were discarded from further analyses, leaving 183 species.

### Model evaluation

We evaluated the predictive performance of the DOMs by comparing species observations to the predicted probability of observing the species (= the combined outcome of occupancy probability and detection probability as the proportion of predicted detections across posterior draws). We assessed whether the models capture temporal trends in occupancy dynamics and how well they capture differences in space (Briscoe et al., 2021). For this, models for the 183 species were refitted to temporal and spatial subsets of data (see below). For 24 species, MCMC diagnostics, assessed as described above, suggested issues in model fitting for at least one data subset. These were discarded from further analyses, leaving 159 species for which models were evaluated. Model fitting issues tended to affect species with a low total number of detections, while the number of routes with detections was similar compared to species for which model fitting was successful. While this could lead to a small bias in the retained species set towards better sampled species, it is unlikely to affect the main results.

To assess the temporal predictive performance, we refitted the models with data from the first 15 years (1995 to 2009) in the same way as described above. We then tested the predictive performance for the following ten years (2010 to 2019) by comparing the number of routes where a species was observed in each year and the number of routes where a species was predicted to be detected. We considered the models to adequately capture the temporal pattern of occupancy dynamics if either (a) the error was comparatively small, which we defined as either the observations being within the 95 % prediction credible interval or the mean absolute percentage error being < 10 %, or (b) the overall trend was captured, which we defined as the Pearson correlation between time series being > 0.5, and additionally (c) if there was no large deviation from expectation, which we defined as the Pearson correlation being not significantly negative and the mean absolute error showing no significant positive trend over the test years.

To assess predictive performance in space, we conducted five-fold spatially-blocked cross validations. We grouped for each species the routes within the 750 km buffer around the detections into spatial blocks and assigned the blocks to five folds using the R package blockCV (Valavi et al., 2019). We tested different hexagon sizes for the spatial blocks and visually checked the resulting fold assignments for each species. We considered a size of 500 km to be suitable across species. We then ran iterations of randomly assigning blocks to folds with the aims of getting an equal number of detections and non-detections in each fold and getting training sets that contain 80 % of the detections, leaving 20 % for the test sets. We then refitted the DOMs in the same way as described above for each fold. To evaluate how well DOMs capture differences in space, we assessed the degree to which routes on which a species was observed have consistently higher proportion of predicted detections across posterior draws than routes where a species was not observed. We quantified this as the mean yearly AUC (area under the receiver operating characteristic (ROC) curve) (Briscoe et al., 2021), using the R package pROC (Robin et al., 2011). We considered an AUC of ≥ 0.7 to indicate acceptable discrimination ability in space (Hosmer Jr et al., 2013).

Taken together, the DOMs for 80 species decently captured temporal trends in occupancy dynamics as well as differences in space. Therefore, we included these 80 species in the final dataset for the impact attribution analysis.

### Attribution

To assess the relative importance of climate and land use change, we used the fitted DOMs of the 80 species with acceptable predictive performance to simulate occupancy dynamics under the following counterfactual scenarios across the study period (Fig. 1b): a) no climate change, but factual land use change, b) no land use change, but factual climate change and c) no climate and no land use change. With this approach, we assumed that DOMs that perform well with factual input data also perform well with counterfactual input data (Mengel et al., 2021), which is conditional on the model structure remaining stable under counterfactual conditions. Since we removed the effects of climate and land use change only from 1995 onwards, the models should rarely encounter input data beyond the range of training data, i.e. extrapolation is limited.

We first assessed how climate and land use change affected the overall trend in occupancy dynamics and identified absolute and relative winners and losers (cf. Langhammer et al., 2024). For each species, we predicted observations as the combined outcome of occupancy probability and detection probability at each route within the 750 km buffer around the species’ observations. This allowed us to calculate meaningful prediction errors based on observations (see below). Then, we calculated the number of occupied routes per species per year by summing the predicted observations from 100 posterior draws, scaled these by the predicted median number of occupied routes in the first year, and fitted a linear model without an intercept to derive the linear trend (Figure S 2). We then assessed the change in slope between factual simulations, which incorporate climate and land use change, and counterfactual simulations. We considered a species to be an *absolute winner* in response to a driver if the number of occupied routes significantly increases in factual simulations and increases less in counterfactual simulations without the driver (Figure 1c). We considered a species to be a *relative winner* if factual simulations suggest a stable or negative trend in the number of occupied routes, but counterfactual simulations suggest a stronger negative trend. Species were categorized as *relative losers* if we found a stable or positive occupancy trend in factual simulations, but a more positive trend in counterfactual simulations. Last, a species was considered to be an *absolute loser* of change if factual simulations showed a significant decrease in occupancy, while counterfactual simulations showed a lower decrease or even increase.

Secondly, we quantified the relative importance of climate and land use change for species’ overall occupancy dynamics while controlling for differences in predictive performance between models. To this end, we defined the relative importance of a driver as the difference between the mean absolute percentage error of factual simulations, that include the driver, and that of counterfactual simulations, that exclude the driver. This implies that a driver is important in proportion to the decrease in model predictive performance when it is not taken into account. Note that this metric of driver importance captures the predictive contribution of a driver rather than causal relationships. We used the median of the posterior distributions as the predicted time series and quantified the mean absolute percentage error with regard to the bird observations with the R package Metrics (Hamner & Frasco, 2018).

To assess whether the effects of climate and land use change were additive, antagonistic or synergistic, we calculated an additivity index as *I* = *imp_CL_* − (*imp_C_* + *imp_L_*), where imp_CL_ = relative importance of the combination of climate and land use change, imp_C_ = relative importance of climate change alone and imp_L_ = relative importance of land use change alone. *I* > 0 indicates synergistic effects of climate and land use change and *I* < 0 antagonistic effects of both drivers. To examine spatial patterns in the relative importance of climate and land use change, we overlaid the breeding, or, for non-migratory species, year-round ranges for the 80 species taken from BirdLife (BirdLife International and Handbook of the Birds of the World, 2022). We then calculated for each grid cell the mean importance of climate or land use change for observed occupancy dynamics across all species whose ranges cover the respective grid cell.

## Results

### Predictive performance

Combining our criteria for adequate temporal and spatial predictive performance resulted in a set of 80 species that were evaluated in the attribution step. Details on the predictive performance of all species for which models were successfully fit are given in Supplemental Material. Of the resulting species, 80 % (64 species) are Passeriformes. Roughly 40 % (34 species) belong to one of four families: Passerellidae (New World sparrows), Tyrannidae (tyrant flycatchers), Cardinalidae (cardinals) and Parulidae (wood-warblers). Roughly one third (24 species) are species of the eastern temperate forests and another third (26 species) occurred only in the western half of the conterminous USA. The retained species do not differ systematically from the species discarded due to poor predictive performance of the models in terms of the total number of detections or the number of routes on which a species was present.

### Factual and counterfactual occupancy trends

Under factual climate and land use, we find a negative trend (linear regression, slope < 0, p < 0.05) in occupancy between 1995 and 2019 for the majority of analysed bird species (45 species, 56 %). The trend in the occupancy dynamics is positive for 31 species (39 %; linear regression, slope > 0, p < 0.05), and insignificant for four species (Figure 2; linear regression, p >= 0.05).

**Figure 2.**
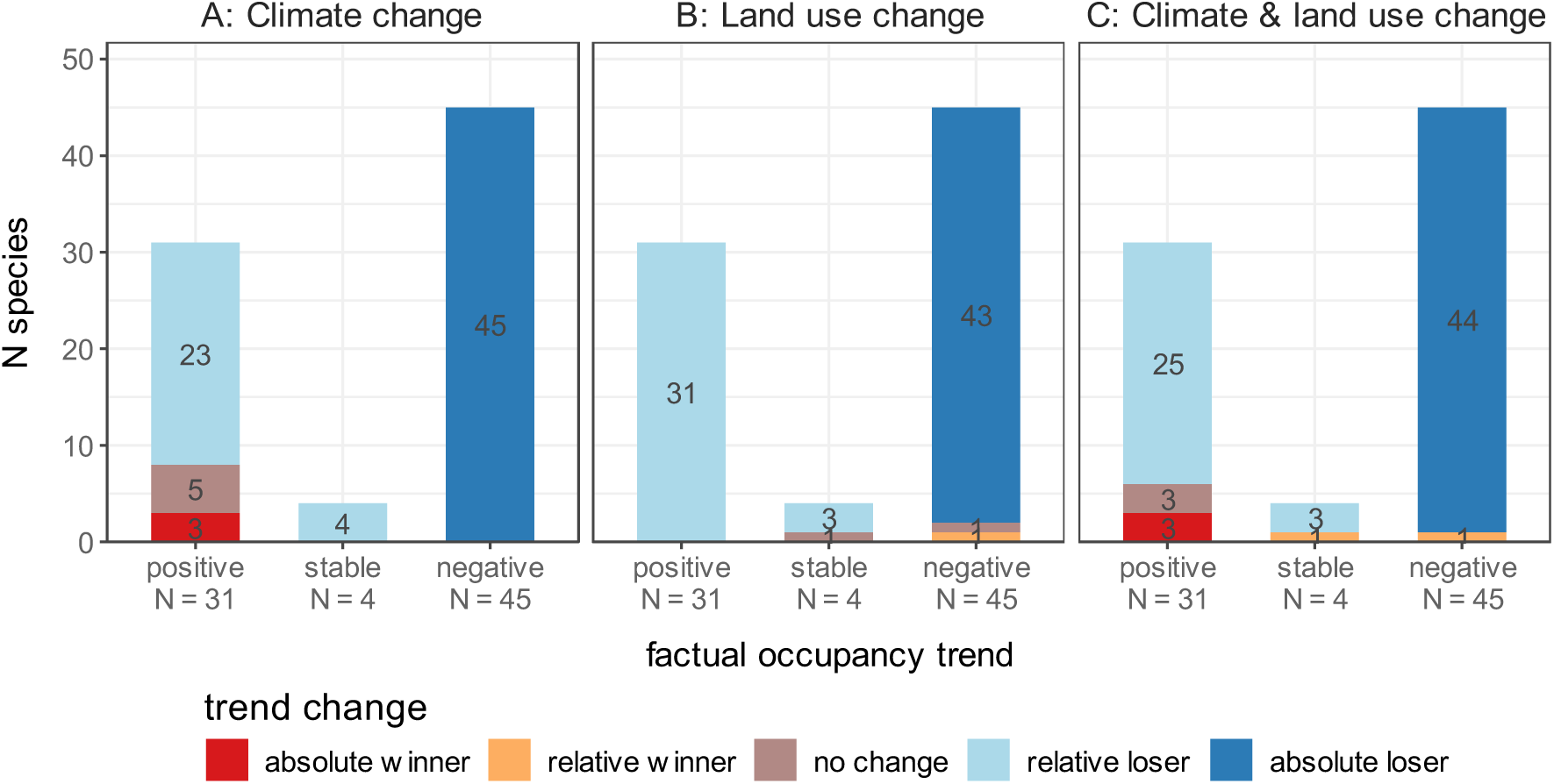
Absolute and relative winners and losers of climate change, land use change and the combination of both in 80 breeding bird species in the conterminous USA over 25 years (1995-2019). Numbers represent number of species. Species are grouped based on the linear trend in occupancy dynamics under observed climate and reconstructed land use (“factual occupancy trend”; x-axis). Winner and loser categories were determined by comparing factual simulations with simulations under counterfactual scenarios of detrended climate (A), constant 1995 land use (B) and both (C). Colours distinguish absolute and relative winners and losers as defined in Figure 1c (*absolute winner*: factual trend > counterfactual trend, factual trend significantly positive; *relative winner*: factual trend > counterfactual trend, factual trend not significantly positive; *relative loser*: factual trend < counterfactual trend, factual trend not significantly negative; *absolute loser*: factual trend < counterfactual trend, factual trend significantly negative). Occupancy time series were simulated with dynamic occupancy models fitted to North American Breeding Bird Survey data (Ziolkowski et al., 2024) and observed climate (Lange et al., 2025) and reconstructed land use data (Volkholz & Ostberg, 2024). Only species for which the models showed acceptable spatio-temporal predictive performance were included.

Our winner and loser classification highlights significant differences in the frequencies of climate change winner and loser species (χ^2^-test, χ^2^ = 94.3, df = 4, p < 0.001). 90 % of the considered bird species are either absolute or relative losers of climate change (Figure 2A). More than half of the 80 species are considered absolute climate change losers (45 species, 56 %), showing a more negative trend under factual than under counterfactual climate. Another 27 species (34 %) can be considered relative losers, meaning that although their occupancy dynamics were either stable or increased under factual climate, we find an even stronger increase in occupancy under the counterfactual climate. In contrast, only three species (4 %) are absolute winners of climate change.

Species classification as winners and losers of land use change also differs significantly from random (χ^2^-test, χ^2^ = 108.1, df = 4, p < 0.001). All species except for three can be considered either absolute losers (43 species, 54 %) or relative losers (34 species, 43 %) of land use change (Figure 2B). One species, Cassin’s sparrow (*Peucaea cassinii*), can be considered a relative winner of land use change.

Comparing factual simulations and simulations with counterfactual climate and constant land use reveals patterns in between those for considering either climate or land use change in isolation (Figure 2C; χ^2^-test, χ^2^ = 91.4, df = 4, p < 0.001).

### Relative importance of different drivers

The relative importance of climate and land use change was quantified as the difference in predictive accuracy between factual and counterfactual simulations. This difference indicates that for 53 species (66 %) climate change was more important for the observed occupancy dynamics than land use change (Figure 3). Across species, the mean relative importance of climate change is 8 % (sd = 4 %, range = 2 % to 15 %), meaning that the mean absolute percentage error on average increases by 8 % under counterfactual climate compared to factual climate. The mean relative importance of land use change is 6 % (sd = 2 %, range = 0 % to 11 %). Note that we consider comparisons of importance values for climate and land use change within and between species to be particularly informative rather than single importance values in isolation. We find the highest importance of climate change for the occupancy dynamics of yellow-bellied sapsucker (*Sphyrapicus varius)* and white-crowned sparrow (*Zonotrichia leucophrys*). Land use change is most important for Bell’s vireo (*Vireo bellii)* and bobolink (*Dolichonyx oryzivorus*).

**Figure 3.**
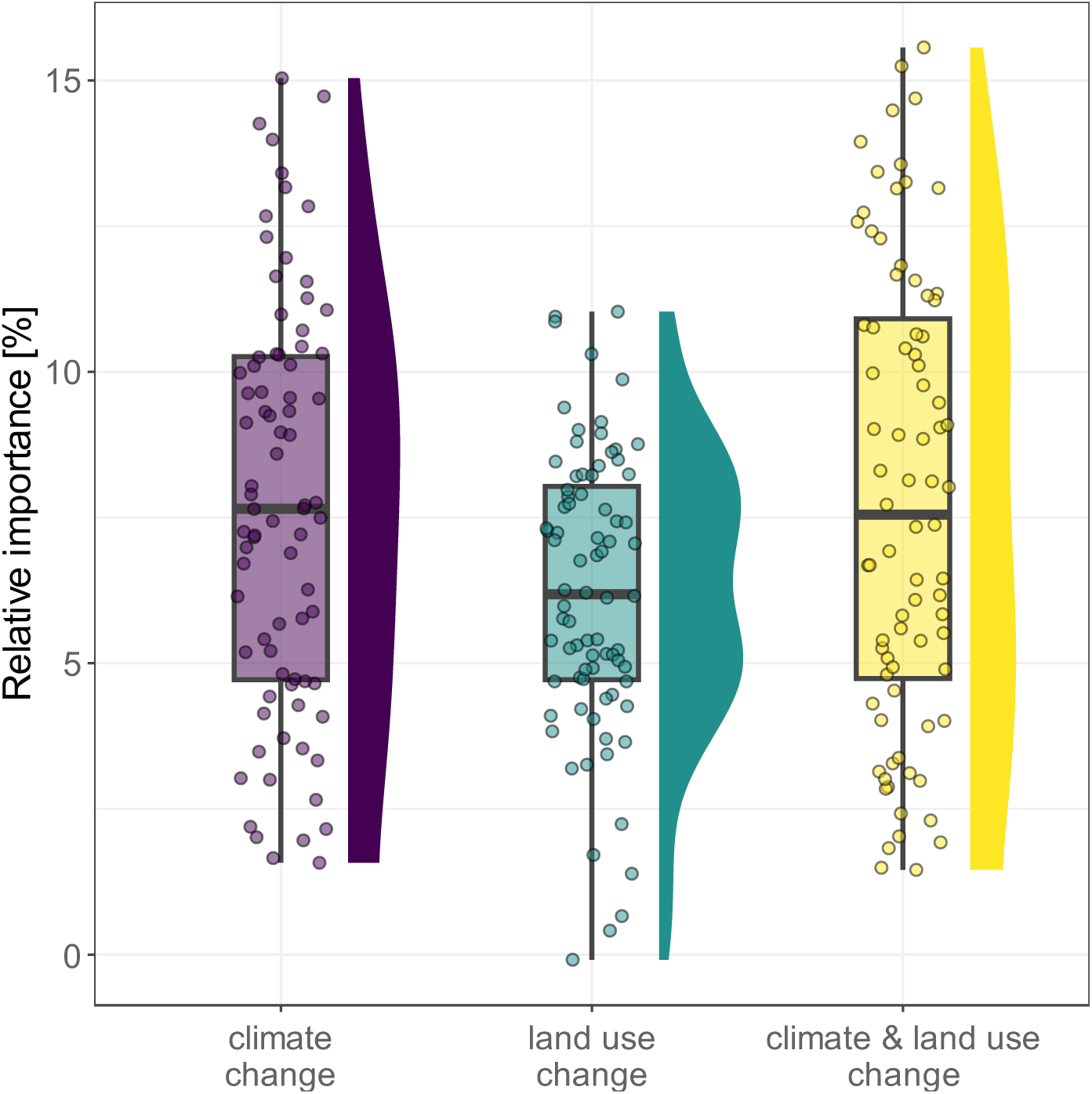
Relative importance of climate change, land use change and the combination of both drivers for observed occupancy dynamics of 80 breeding bird species in the conterminous USA over 25 years (1995 – 2019). Relative importance was quantified as the difference between the mean absolute percentage error of the factual simulation, i.e. simulated occupancy time series for observed climate and reconstructed land use, and the counterfactual scenarios (left: no climate change, middle: no land use change, right: no climate and no land use change). Time series were simulated with dynamic occupancy models fitted with observed climate (Lange et al., 2025) and reconstructed land use data (Volkholz & Ostberg, 2024) and bird observations from the North American Breeding Bird Survey (Ziolkowski et al., 2024). Each dot shows the difference in mean absolute percentage errors for one species, boxplots depict summary statistics, density curves show the distribution of the data.

At the species level, we find that a single species can be a winner of one driver of change, but a loser of another. Based on our analysis, Bell’s vireo, for example, is an absolute winner of climate change, but a relative loser of land use change, with land use change being a more important driver for the species’ occupancy dynamics than climate change (Figure 4).

**Figure 4.**
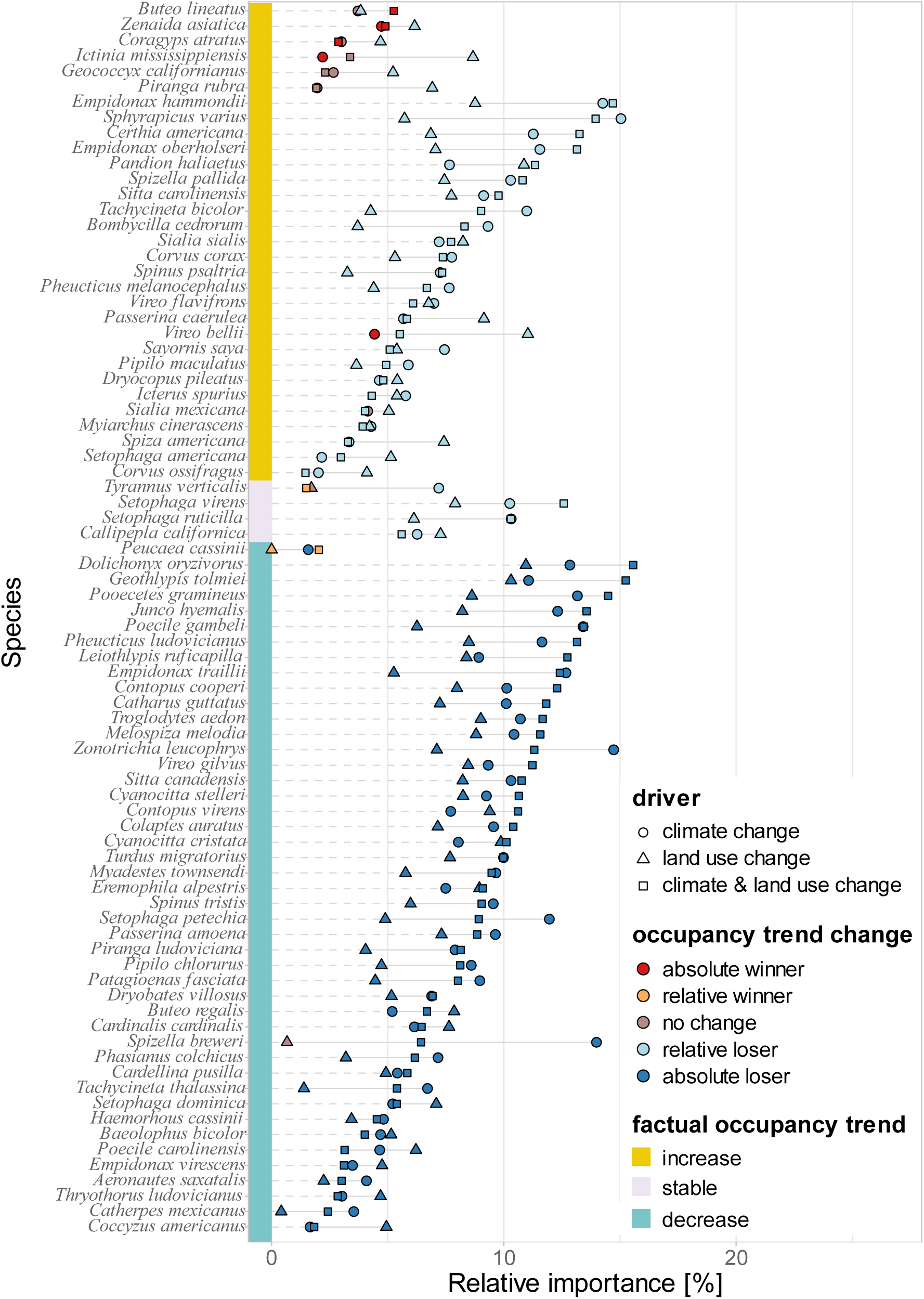
Species-level relative importance of climate change, land use change and the combination of both drivers for observed occupancy dynamics of 80 breeding bird species in the conterminous USA over 25 years (1995 – 2019). Occupancy trend change categories are defined in Figure 1c (absolute winner: factual trend > counterfactual trend, factual trend significantly positive; relative winner: factual trend > counterfactual trend, factual trend not significantly positive; relative loser: factual trend < counterfactual trend, factual trend not significantly negative; absolute loser: factual trend < counterfactual trend, factual trend significantly negative). Relative importance was quantified as the difference between the mean absolute percentage error of the simulated occupancy time series for the factual simulation (i.e. observed climate and reconstructed land use) and the simulated time series for counterfactual scenarios with detrended climate and/ or constant 1995 land use. The coloured bar on the left depicts the factual linear trend in occupancy and corresponds to the x-axis in Figure 2. Time series were simulated with dynamic occupancy models fitted with observed climate (Lange et al., 2025) and reconstructed land use data (Volkholz & Ostberg, 2024) and bird observations from the North American Breeding Bird Survey (Ziolkowski et al., 2024).

The combination of climate and land use change is more important for 31 species (39 %) than climate change and land use change alone (Figure 4). For all species, except for Cassin’s sparrow (*Peucaea cassinii*), we find antagonistic effects of climate and land use change (Figure S 3; additivity index: mean: – 6 %, median: -6 %, range: -11% to -2%). This means that the effects of climate and land use change may partially balance each other out. For Cassin’s sparrow our results indicate slightly synergistic effects of both drivers (additivity index + 0.5%).

### Spatial patterns

Overlaying the breeding, or, for non-migrants, the year-round ranges (BirdLife International and Handbook of the Birds of the World, 2022) indicates that climate change is particularly affecting occupancy dynamics of species that occur in the western part of the conterminous USA, especially at higher elevations, as well as in the northeastern parts and in the Appalachian Mountains (Figure 5A). These species are mainly relative or absolute climate change losers (Figure S 5). Absolute climate change winners tend to have their ranges in the southern USA and in parts of the Great Plains. Land use change particularly affected occupancy dynamics of species that occur in the eastern conterminous USA (Figure 5B), with no particular spatial patterns in the ranges of relative and absolute losers (Figure S 6, compare to Figure S 4). In large parts of the conterminous USA, the combination of climate and land use change was the most important driver of occupancy change at the range level across bird communities, particularly in the Northern Forests, the Appalachian Mountains and the western Great Plains (Figure 5C). For bird communities in the western regions, however, climate change alone was the dominating driver, except for the coastal areas. Land use change was the most important driver for bird communities in the southeast from Texas to North Carolina.

**Figure 5.**
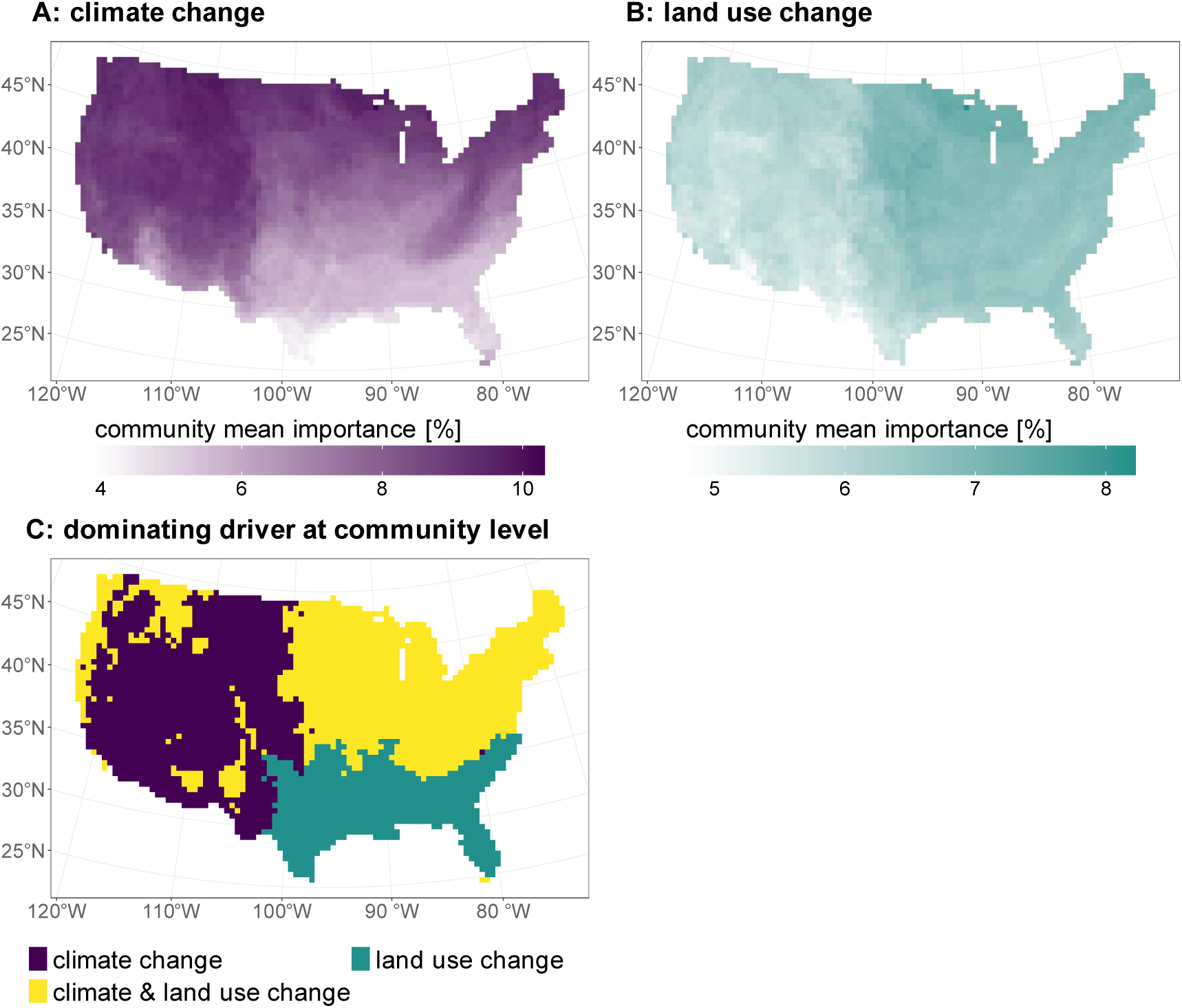
Community mean relative importance of climate change (A) and land use change (B) as well as driver dominance (C) quantified at the range level of 80 species. The darker the colour in A and B, the more strongly are the species in the local community affected by climate (A) or land use change (B) throughout their range within the conterminous USA. We overlaid the breeding, or, for non-migratory species, year-round ranges for 80 species of North American breeding birds, taken from BirdLife (BirdLife International and Handbook of the Birds of the World, 2022). We then calculated for each grid cell the mean importance of climate change, land use change and of both drivers combined for the occupancy dynamics across all species which ranges cover the respective grid cell. Note the different ranges of the colour scales in A and B. C shows the driver with the largest community mean relative importance. Relative importance for single species was quantified as the difference between the mean absolute percentage error of the simulated occupancy time series for observed climate and reconstructed land use and the simulated time series for counterfactual scenarios with detrended climate and/ or constant 1995 land use. Time series were simulated with dynamic occupancy models fitted with observed climate and reconstructed land use data (Lange et al., 2025; Volkholz & Ostberg, 2024) and bird observations from the North American Breeding Bird Survey (Ziolkowski et al., 2024).

## Discussion

In this study, we applied an impact attribution framework to disentangle the relative contributions of climate and land use change to recent occupancy dynamics of North American breeding birds. Based on dynamic occupancy models, which describe occupancy change explicitly via colonisation and local extinction and which we fitted to long-term monitoring data, we show that most species were relative or absolute losers from climate change and land use change, and that climate change emerged as overall more important driver of changes in species occupancy since the mid-1990s. This result is consistent with a slowing down of land use change in many regions of the USA (excluding suburbanization) and an accelerating impact of climate change during this period. Climate change impacts were particularly important for species occurring in the western USA, while land use change was more influential on species in the southeast. Notably, the effects of climate and land use change have partially offset each other rather than being additive for almost all species. Together, our results provide a robust, data-driven assessment how different global change drivers have affected recent biodiversity dynamics at continental scale.

Our results suggest that for many bird species climate change has affected occupancy dynamics over the last 25 years more strongly than land use change, while previous assessments consistently reported land use change as more important driver explaining historic biodiversity change (IPBES, 2019). One reason for this could be a higher exposure to climate change during the study period. While climate change in general intensified from the 1980s onwards, with global mean temperature increasing from 1995 to 2019 by approximately 0.7 °C (Gulev et al., 2021), major land conversions in the conterminous USA took place before our study period. Cropland areas in North America stabilized by the 1950s and many temperate biomes had been converted by 1990 (Millennium Ecosystem Assessment, 2005). Thus, between 1995 and 2019, breeding birds in many regions of the conterminous USA were likely exposed to higher levels of climate change compared to broad-scale land use changes (cf. Figure S 8). It is perhaps not surprising therefore, that we found that climate change had a greater impact than land use change on occupancy of many species across our study period, although land use impacts were still pronounced. If we had conducted our study prior to this period, we would expect that land use change would have emerged as the more important driver. Although beyond the scope of this attribution study and hampered by availability of high-quality long-term biodiversity data, future work should aim to incorporate broader time periods to capture a range of dynamics of key drivers.

Another reason for the larger impact of climate change compared to land use change could be that the species we modelled are overall more sensitive to changes in climate than to changes in land use. While disentangling this is beyond the scope of our analysis, notably, the top species for each driver show trait-coherent patterns: top land-use-change-impacted species are habitat specialists (grassland obligates like bobolink, riparian scrub obligates like Bell’s vireo), while top climate-change-impacted species are thermally constrained (montane/boreal species like yellow-bellied sapsucker and white-crowned sparrow), consistent with trait-based vulnerability frameworks (Jiguet et al., 2010; Pearce-Higgins et al., 2015).

Alternatively, the larger importance of climate change compared to land use change could result from climate change impacts being better captured in the models than land use change impacts. While climate change is acting at large spatial scales, important processes associated with land use change, such as habitat fragmentation and agricultural intensification, may act at scales too small to be adequately captured at the 0.5° resolution applied here. While our land-use variables reflect changes in the proportion of the cell covered by each land cover type (Hurtt et al., 2020), relating these to occupancy at the route level may obscure important patterns (Ankori-Karlinsky et al., 2022). Additionally, the available land use variables may be less suited to capture important aspects related to bird ecology than the available climate variables. For example, the land use data distinguish primary and secondary forests, while birds might select a particular forest type such as evergreen or deciduous. For future attribution studies, it would therefore be desirable to compile land use and species occurrence data with a higher spatial resolution and fit models at more local scales to capture such patterns. In addition, such efforts should focus on land use related variables that are particularly relevant to species’ ecology.

According to our attribution analyses, the majority (55 %) of the 80 bird species studied in the conterminous USA can be considered absolute losers of recent climate and land use change with widespread declines in occupied area. This is consistent with severe continent-wide abundance declines found across North American bird species during recent decades (Rosenberg et al., 2019; Johnston et al., 2025). The estimated higher impact of climate change over land use change for the majority of the studied species conflicts with the prevailing view that land use change is still the dominant driver of biodiversity change across the globe (Sala et al., 2000; IPBES, 2019). However, a recent attribution study by Kotz et al. (2025) suggests that the relative importance of these drivers is shifting over time. The authors found that the impact of human pressure, including land use change, on bird abundance change has stabilized from 2000 onwards, while the impact of climate change intensified since the 1980s (Kotz et al., 2025). This is consistent with our finding that climate change has emerged as a more important driver of occupancy changes for bird species.

At regional scales, the relative importance of climate and land use change depends on environmental context and on the aspect of biodiversity considered (IPBES, 2019). Previous studies reported increases in annual mean temperature between 1980 and 2015 particularly in the western half of the conterminous USA (Currie & Venne, 2017; Hicke et al., 2022) and found that poleward shifts in peak bird abundances were most pronounced in this region (Martins et al., 2024). This is consistent with our result that climate change is particularly important for species that occur in the western USA. There, it was especially the annual temperature range that changed throughout our study period (Figure S 8), which suggests that changes in seasonal temperature variation may be important for understanding climate change effects on bird occupancy dynamics. Although northern range limits remained relatively stable over the last decades and the centres of distributions shifted predominantly westwards (Currie & Venne, 2017; Zurell et al., 2024), within their ranges species occupy sites that are less exposed to warming (Currie & Venne, 2017). Such redistributions can precede range shifts (Billman et al., 2025; Kelly & Goulden, 2008), and suggest, consistent with our results, that climate change is already a key driver of occupancy dynamics.

Ecological models have rarely been combined with attribution approaches from climate science to assess biodiversity responses to global change (Gonzalez et al., 2023; Dudney et al., 2025, examples are Dudney et al., 2021; Erazo et al., 2024; Hari et al., 2026; Kotz et al., 2025). The explicit representation of colonisation and local extinction allows DOMs to capture non-equilibrium responses to environmental change, which is particularly relevant for attribution of occupancy change over short time periods. Our analyses build on standardized long-term monitoring data and focus on consistently surveyed locations to reduce biases, which makes them well-suited for attribution studies (Schrodt et al., 2025; Ziolkowski et al., 2024). Importantly, we ensured that model simulations agree with observed changes (Mengel et al., 2021; Frieler et al., 2024; Gonzalez et al., 2023) by only retaining models with acceptable predictive performance in our retrospective counterfactual modelling (Dudney et al., 2025). Taken together, this framework provides a general and scalable approach for attributing biodiversity change to multiple interacting drivers using observational data. More broadly, assessing responses at the species level is important because species differ in their sensitivity to global change drivers (Dawson et al., 2011; McGill et al., 2015; Williams et al., 2008). As most ecological models at the species level need to be calibrated with data (Briscoe et al., 2019), data-driven impact attribution represents an important step towards operationalizing attribution in biodiversity science.

Spatial and temporal predictive performance of the DOMs was acceptable for only 80 of the initially considered 192 bird species, despite having excluded species beforehand for which we expected occupancy dynamics to be challenging to model (particularly rarer species) and using a subset of the BBS data that provides consistent observation time series with minimal sampling variability. This underscores the importance of evaluating and improving modelling pipelines to better characterise species’ responses to varying environmental conditions, which is a prerequisite for attributing biodiversity change to drivers. Accurate models of ecological responses are lacking for the vast majority of species on Earth (Urban et al., 2016, 2022; Zurell et al., 2026). Ecological systems are complex and often affected by several interacting drivers that mediate species responses. Additionally, ecological responses to global change are often transient and lag behind drivers owing to long generation times and buffering mechanisms (Essl et al., 2024; Schrodt et al., 2025). While the intermediate complexity of DOMs makes it feasible to fit them for many species if data of repeated surveys are available (Kelleher et al., 2025), alternative approaches that incorporate lower-level processes leading to occupancy change, such as demography, physiology, species interactions, (local) adaptation and dispersal capability (Urban et al., 2016; Briscoe et al., 2019) can provide a more mechanistic understanding of how environmental change translates into changes in occupancy and population dynamics. However, realistically modelling these processes for many species remains challenging due to a lack of relevant data and technical barriers regarding model calibration (Briscoe et al., 2019; Urban et al., 2016; Zurell et al., 2016).

One limitation to our study is that we cannot rule out the influence of drivers other than climate and land use change. Factors such as large-scale changes in agricultural intensification, including increased use of neonicotinoids (Li et al., 2020), forestry (Akresh et al., 2023), pollution (Hallmann et al., 2014; Nygård et al., 2019) or changes regarding direct human interactions with birds, e.g. via supplementary feeding (Greig et al., 2017) may have increased in parallel to warming temperature. Thus, a part of what we attributed to climate change may in fact be caused by factors covarying with global warming. Disentangling the impacts of additional drivers is challenging due to limited availability of driver data with sufficient spatio-temporal coverage. As new data become available, future attribution studies could additionally integrate a broader range of drivers to provide a more complete picture of the causes of biodiversity change (Schrodt et al., 2025; Thomas et al., 2026), which may also improve our ability to adequately model dynamics across a broader range of species. To capture the full extent of climate change impacts, future studies could incorporate variables reflecting exposure to climatic extremes, since these were recently found to be particularly important for driving bird abundance changes (Kotz et al., 2025). Excluding rarer species, for which it is challenging to obtain informative models (Zhang et al., 2020), may limit the extent to which our results reflect impacts of change across all breeding bird species in the USA. Since rarer species may be particularly sensitive to changing climatic conditions (Vincent et al., 2020), our results for the set of species we considered could represent conservative estimates of the impacts of climate change across breeding bird species in the USA. With regard to migratory species, our analyses likely miss climate and land use change impacts that act outside of the breeding range on wintering grounds or along flyways. Ultimately, studies should aim at incorporating all areas that are relevant during the annual cycle of species. To aid this endeavour, it is particularly important to continue and expand standardized long-term monitoring efforts (Schrodt et al., 2025; Thomas et al., 2026). A further extension of our approach would be to assess global change impacts at finer spatial scales, which may reveal important spatial variation in driver impacts (Martins et al., 2024; Johnston et al., 2025) and represents a promising direction for future work. While we focus here on aggregated patterns across conterminous USA, our approach is scalable to other spatial resolutions given sufficient data availability.

In summary, impact attribution based on counterfactual scenarios provides a powerful framework to disentangle the impacts of multiple drivers on biodiversity change. By linking observed trends to specific causes, such approaches can improve our understanding of biodiversity dynamics (McGill et al., 2015; Dudney et al., 2025) and provide the basis for robustly projecting future biodiversity change under alternative global change scenarios (Schrodt et al., 2025). Such projections can support policy-making related to meeting international biodiversity targets as well as targets regarding ecosystem functioning, the provision of ecosystem services and risks associated with the spread of diseases (IPBES, 2019; Gonzalez et al., 2023; Zurell et al., 2026). Our results suggest that over the last 30 years, climate change has already been a more important driver of bird occupancy change in the conterminous USA than land use change, in particular in the west. This underscores the need to consider effects of climate change in nature conservation planning, in addition to improving the availability of habitat under current environmental conditions (Langhammer et al., 2024), and to strengthen climate change mitigation efforts.

## Acknowledgements

We acknowledge funding support by the Deutsche Forschungsgemeinschaft (DFG, German Research Foundation), project 518316503 (KS, DZ). We thank Matthias Mengel and Sven Willner for support on generating counterfactual climate data. We also thank everyone who contributed to the North American Breeding Bird Survey. This work used resources of the Deutsches Klimarechenzentrum (DKRZ) granted by its Scientific Steering Committee (WLA) under project bb0820.

