## Supplemental Material for "North American bird occupancy dynamics attributed to climate and land use change"

### Table of Contents

Supplementary information ... 3
Dynamic occupancy models (DOMs) ... 3
Variable selection ... 3
Assumptions underlying the DOMs ... 3
Results – predictive performance ... 4
Supplementary tables ... 5
Route selection criteria ... 5
Supplementary figures ... 6
Map of the selected routes ... 6
Example illustrating occupancy trend calculation ... 7
Boxplot additivity index ... 8
Map of range-level species richness of the assessed species ... 9
Maps of range-level distribution of winners and losers of climate and land use change ... 10
Maps of change in model covariates from 1995 to 2019 ... 13
References ... 16

### Supplementary information

#### Dynamic occupancy models (DOMs)

##### Variable selection

To assess the impacts of climate and land use change across species, we used the same set of covariates for each species. We selected variables that cover a substantial amount of environmental variation across the conterminous USA and that are not substantially correlated. We conducted a principal component analysis (PCA) of the candidate covariates (Briscoe et al., 2021; König et al., 2021) with the R package ade (Dray & Dufour, 2007) and quantified the importance of a variable for capturing environmental variation as the variance captured by the principal axis on which the variable had its highest loading, multiplied by the loading of the variable on that axis. We then calculated Spearman's rank correlations for each pair of variables. If $|\rho| > 0.7$, we removed the  less important variable (Dormann et al., 2013), according to the PCA-based assessment, with the exception that we prioritized annual mean temperature given its general relevance to climate change. This resulted in a selection of 15 variables, ten related to climate: annual mean temperature, mean diurnal temperature range, annual temperature range, temperature isothermality, precipitation seasonality, precipitation of driest month and mean spring, summer, autumn and winter precipitation; and five variables related to land use: primary non-forest, secondary non-forest, managed pastures, urban areas and annual crops.

##### Assumptions underlying the DOMs

Since DOMs require data from repeated surveys to estimate detection probabilities, we considered each route as a single location and five sections along a route, consisting of ten stops each, as replicate observations. By doing so we defined occupancy at the route level, i.e. as the presence of a species along a BBS route during one breeding season (Royle & Kéry, 2007; Jarzyna & Jetz, 2017; Doser et al., 2023). We assumed that a species present on a route is available for detection at any of the sections. We considered this a reasonable assumption for the majority of the selected species, although it depends on each species' individual home range size, habitat configuration and density compared to the route length of approximately 40 km and the route's spatial configuration. We further assumed that there are no false-positive detections, that species occurrence is independent across routes given the distance of at least 100 km and that detection probability is independent across route sections given the standardized protocol of the BBS. Although we recognize that these assumptions are unlikely to hold for all species at all times, they are likely to hold for most of the species in our dataset, and making these decisions is necessary to create tractable, yet realistic models.

#### Results – predictive performance

Across the 159 species for which models were successfully fit to all subsets of data, 81 (51%) met our criteria for adequate temporal predictive performance. The mean absolute percentage error of the predictions was less than 10 % for 56 species, and for 16 additional species the 95 % credible interval covered the observations. For 12 additional species, the observed and predicted time series were correlated with Pearson's $r > 0.5$. Of these 84 species, three species were discarded due to either a significant positive trend in mean absolute error or a significant negative correlation between observed and predicted time series. In contrast, almost all (n=157, 99%) species met our criteria for adequate spatial predictive performance. Across these species, AUC values ranged from a 0.72 to 0.98, with a mean of 0.88 indicating an excellent discrimination ability of the models on average across species (Hosmer Jr et al., 2013).

### Supplementary tables

#### Route selection criteria

**Table S1. Selection of routes from the North American Breeding Bird Survey (Ziolkowski et al., 2024) used to fit dynamic occupancy models.**

| Route selection rationale | Selection criteria | Number of routes |
|---|---|---|
| consider only high-quality observation time series | BBS RunType = 1 (route randomly established, survey follows the official BBS sampling protocol, conducted on a suitable date and time under suitable weather conditions) | 3932 |
| | full spatial information available[1] | 3229 |
| | surveyed in 20 of 25 years (i.e. 80%) between 1995 and 2019, including 1995 and 2019 | 986 |
| reduce bias in spatial coverage | 100 km minimum distance of route centroids | 632 |
| | maximum 30 routes per Bird Conservation Region[2] | 539 |

[1] Patuxent Wildlife Research Center, 1999

[2] Jarzyna & Jetz, 2017

### Supplementary figures

#### Map of the selected routes

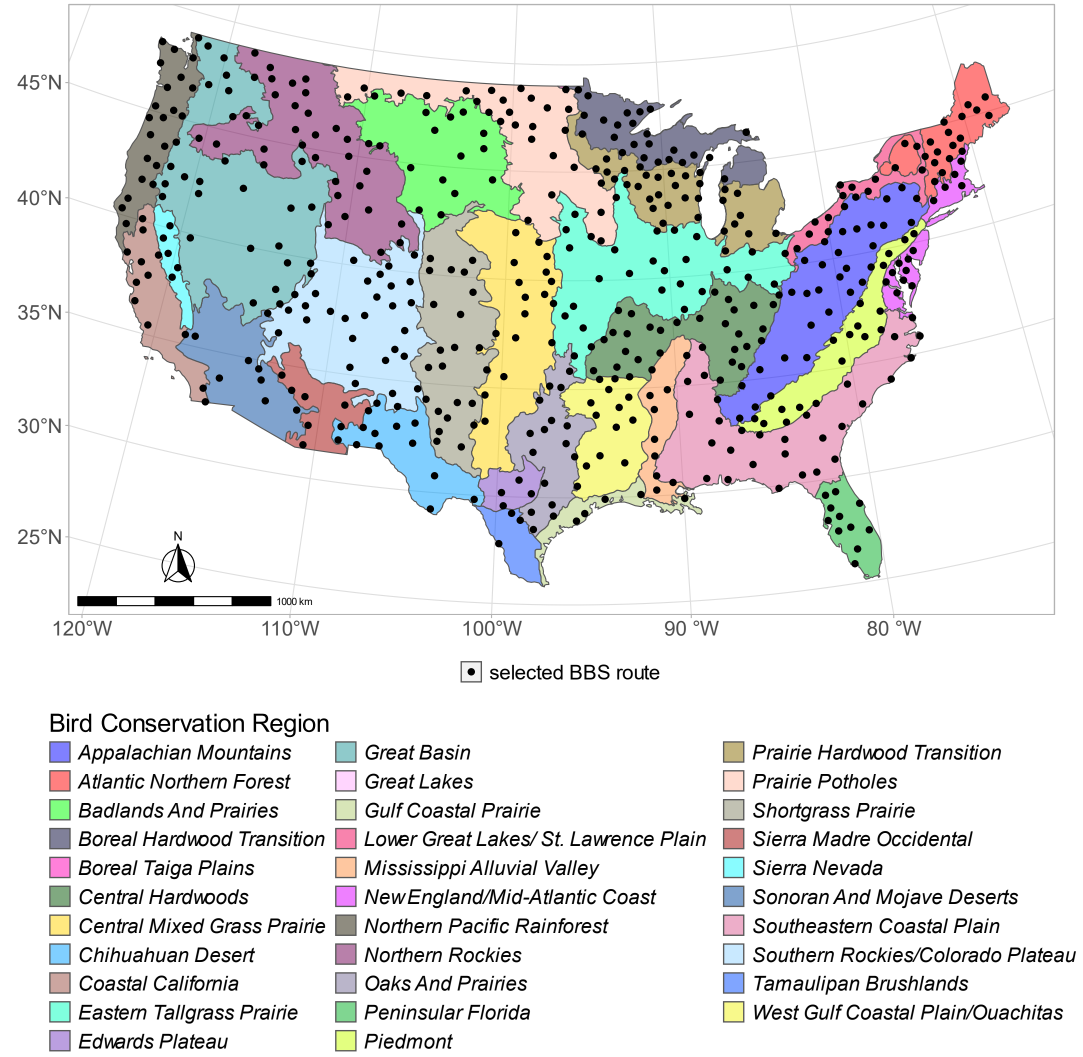


**Figure S 1. Locations of the 539 routes of the North American Breeding Bird Survey (Ziolkowski et al., 2024) selected for fitting dynamics occupancy models and Bird Conservation Regions (Bird Studies Canada and NABCI, 2014) in the conterminous USA.**

#### Example illustrating occupancy trend calculation

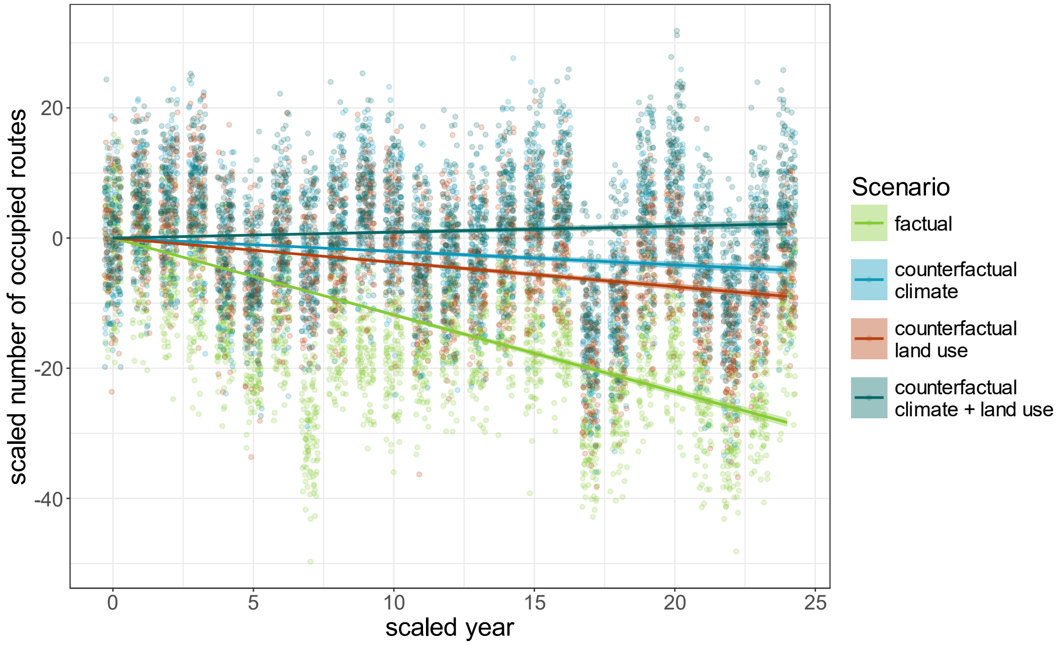


**Figure S 2. Example of how occupancy trends were calculated to determine absolute and relative winners and losers of climate and land use change.** Points show 100 draws of the posterior distributions of the number of occupied routes in each year, simulated with a dynamic occupancy model fitted to bird observations (Ziolkowski et al., 2024) and factual climate and land use data (Lange et al., 2025; Volkholz & Ostberg, 2024) and scaled with the median of the first year. Lines show the linear models, fitted without an intercept, including the standard error. To determine absolute and relative winners and losers of change, we compared the occupancy trends across scenarios based on the slopes of the linear models. Factual scenario = observational climate and land use between 1995 and 2019, including the effect of climate and land use change; counterfactual climate = no climate change since 1995, but factual land use change; counterfactual land use = land use kept constant at the level of 1995, but factual climate change; counterfactual climate + land = no climate change since 1995 and constant land use since 1995. This example shows predictions for the bobolink (*Dolichonyx oryzivorus*).

#### Boxplot additivity index

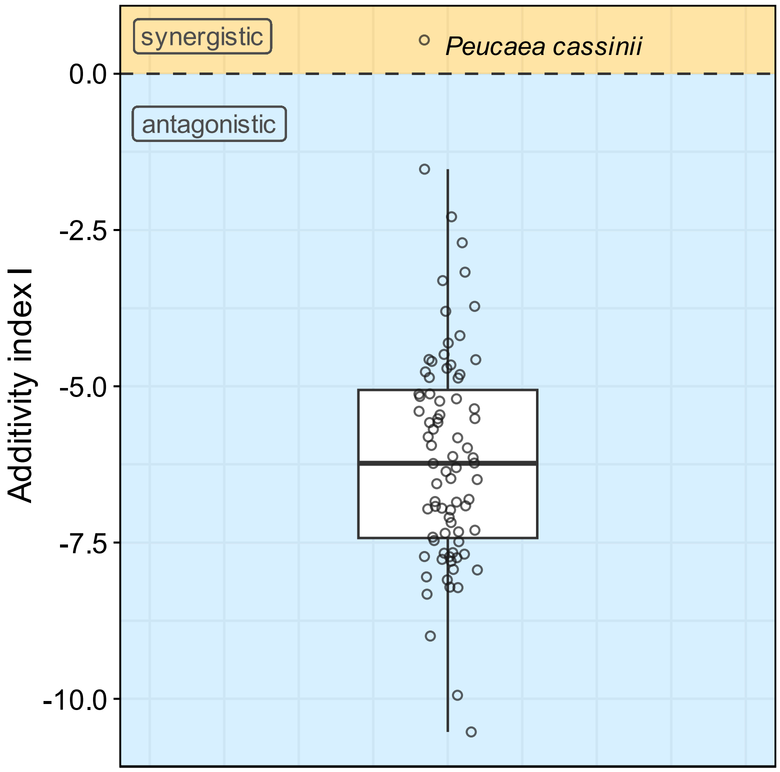


**Figure S 3. Additivity index describing whether the combined effects of climate and land use change on occupancy dynamics of 80 breeding bird species in the conterminous USA over 25 years (1995 – 2019) were antagonistic (I < 0) or synergistic (I > 0).** The index was calculated as $I = imp_{CL} - (imp_C + imp_L)$, where $imp_{CL}$ = relative importance of the combination of climate and land use change, $imp_C$ = relative importance of climate change alone and $imp_L$ = relative importance of land use change alone. Relative importance was quantified as the difference between the mean absolute percentage error of a factual simulation of occupancy dynamics, i.e. simulated occupancy time series for observed climate and reconstructed land use, and the mean absolute percentage error of counterfactual scenarios. Time series were simulated with dynamic occupancy models fitted with observed climate (Lange et al., 2025) and reconstructed land use data (Volkholz & Ostberg, 2024) and bird observations from the North American Breeding Bird Survey (Ziolkowski et al., 2024).

#### Map of range-level species richness of the assessed species

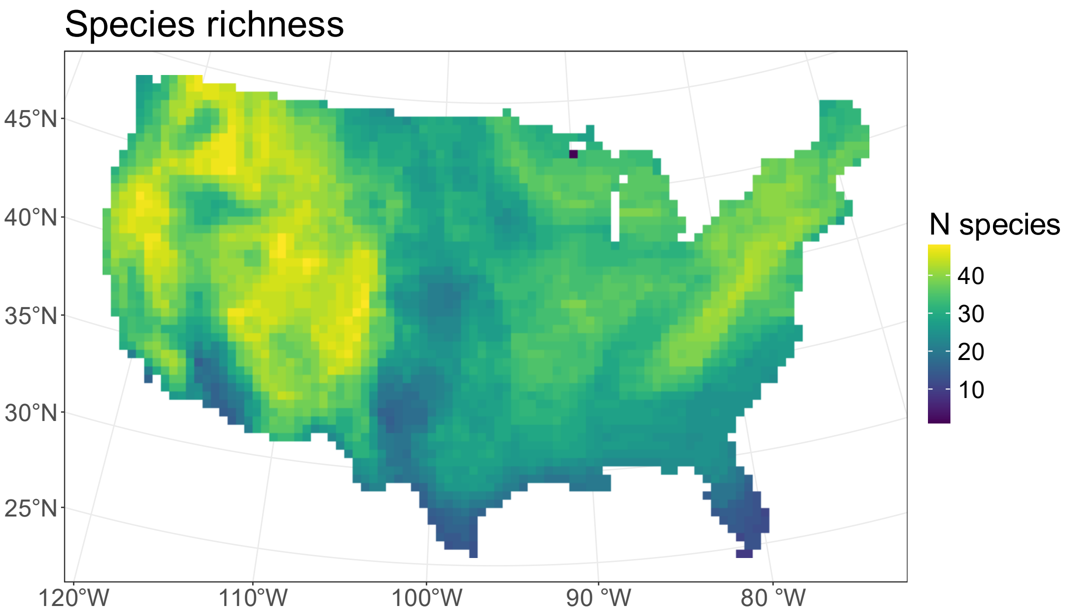


**Figure S 4. Species richness across the conterminous USA of the 80 species for which impacts of climate and land use change on occupancy dynamics were assessed in the attribution step.** We overlaid the breeding, or, for non-migratory species, year-round ranges from BirdLife (BirdLife International and Handbook of the Birds of the World, 2022) and summed for each grid cell the number of species which ranges cover the respective cell.

#### Maps of range-level distribution of winners and losers of climate and land use change

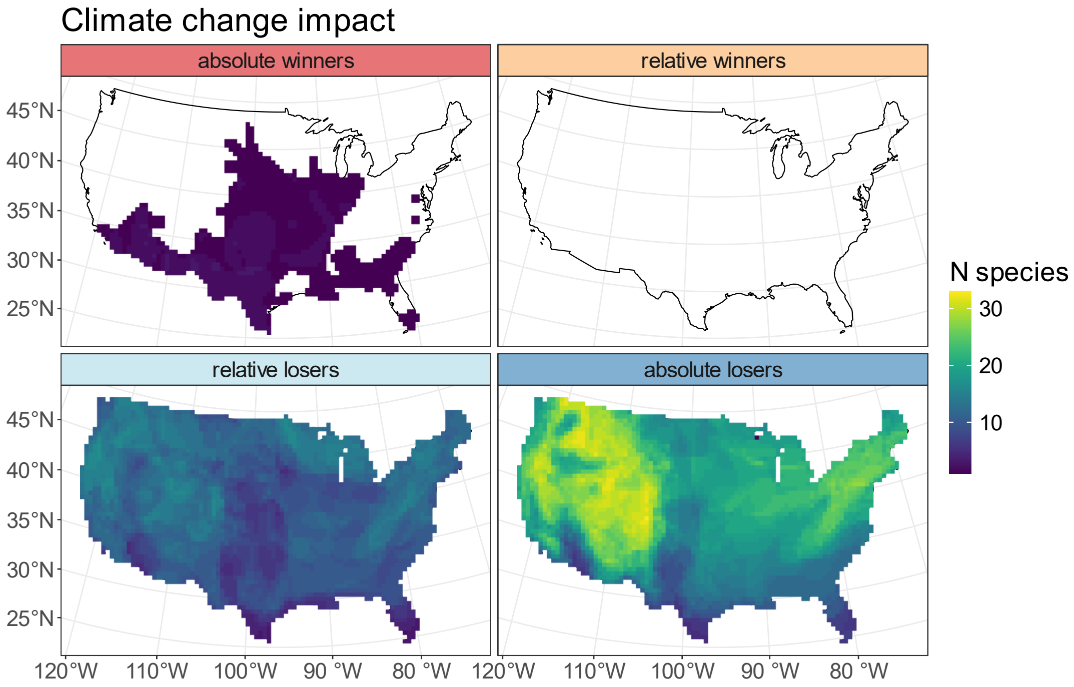


**Figure S 5. Distribution of species classified as relative or absolute climate change winners or losers.** We overlaid the breeding, or, for non-migratory species, year-round ranges for 80 species of North American breeding birds from BirdLife (BirdLife International and Handbook of the Birds of the World, 2022). We then summed for each grid cell the number of species in each impact category which ranges cover the respective grid cell. See **Error! Reference source not found.** on how impact categories were determined.

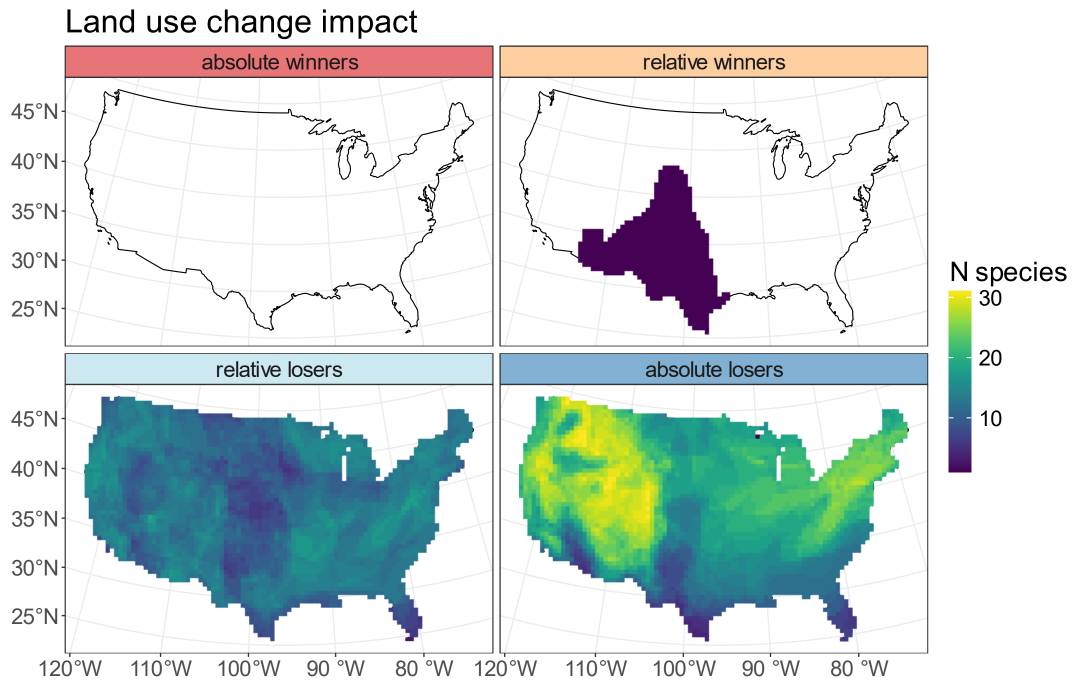


**Figure S 6. Distribution of species classified as relative or absolute land use change winners or losers.** We overlaid the breeding, or, for non-migratory species, year-round ranges for 80 species of North American breeding birds from BirdLife (BirdLife International and Handbook of the Birds of the World, 2022). We then summed for each grid cell the number of species in each impact category which ranges cover the respective grid cell. See Error! Reference source not found. on how the impact categories were determined.

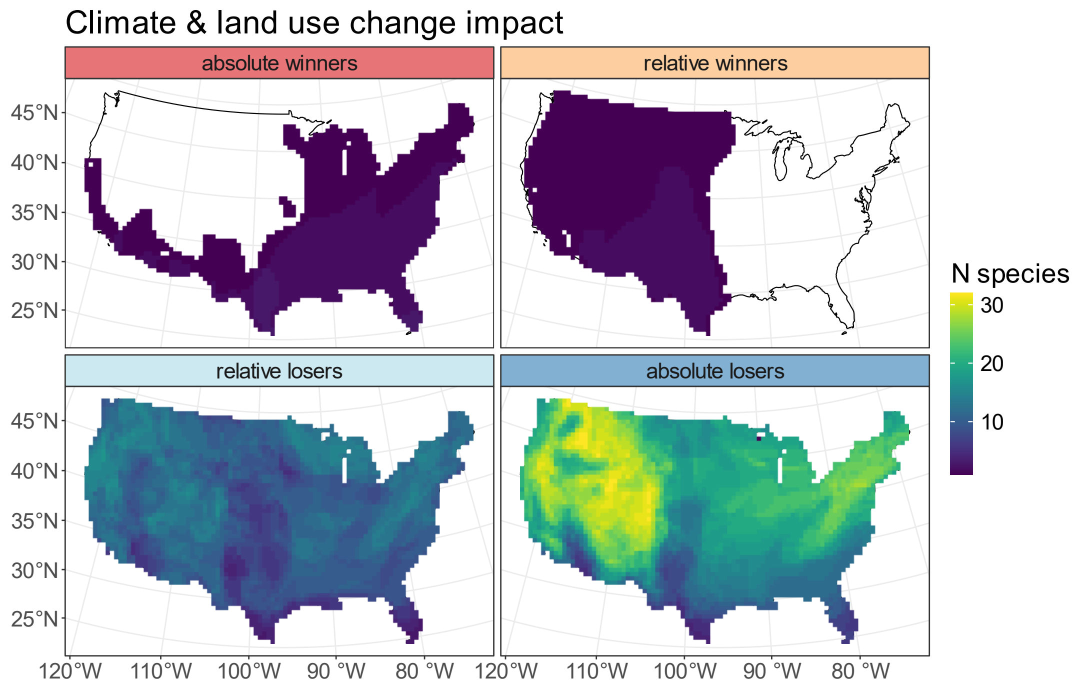


**Figure S 7. Distribution of species classified as relative or absolute winners or losers of the combination of climate and land use change.** We overlaid the breeding, or, for non-migratory species, year-round ranges for 80 species of North American breeding birds from BirdLife (BirdLife International and Handbook of the Birds of the World, 2022). We then summed for each grid cell the number of species in each impact category which ranges cover the respective grid cell. See Error! Reference source not found. on how the impact categories were determined.

#### Maps of change in model covariates from 1995 to 2019

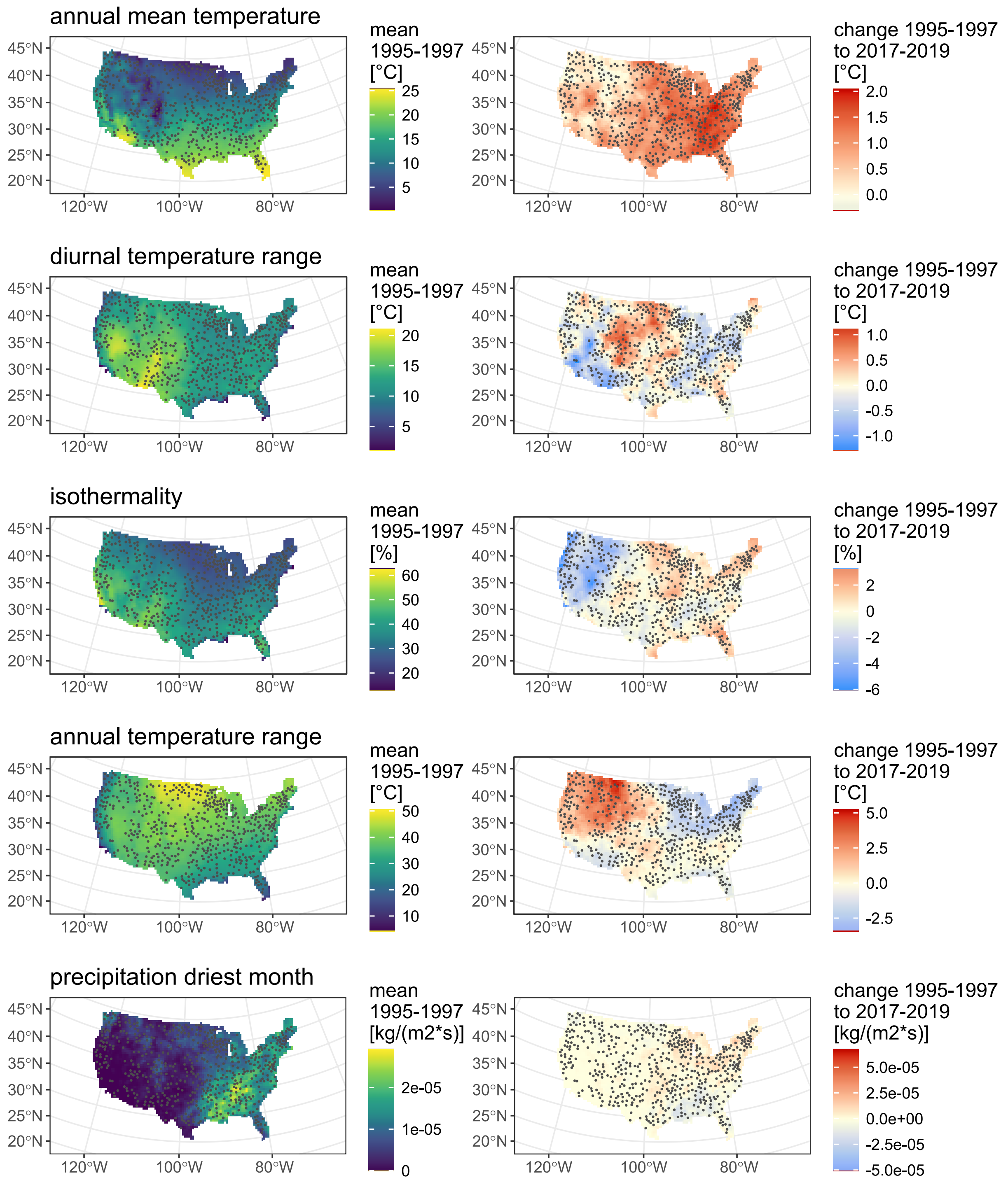


**Figure S 8 (1 of 3). Change in environmental variables from the start to the end of the study period (1995 – 2019) in the conterminous United States.** The left column shows the mean values across the first three years. The right column shows the change from the first to the last three years of the study period as the difference between the mean value of 1995-1997 and the mean value of 2017-2019. The variables refer to observed climate (Lange et al., 2025) and reconstructed land use (Volkholz & Ostberg, 2024) and were used to fit dynamic occupancy models for breeding bird species. Note the different units and ranges of the colour scales. Dots show the location of 539 routes of the North American Breeding Bird Survey (Ziolkowski et al., 2024) used to fit the models.

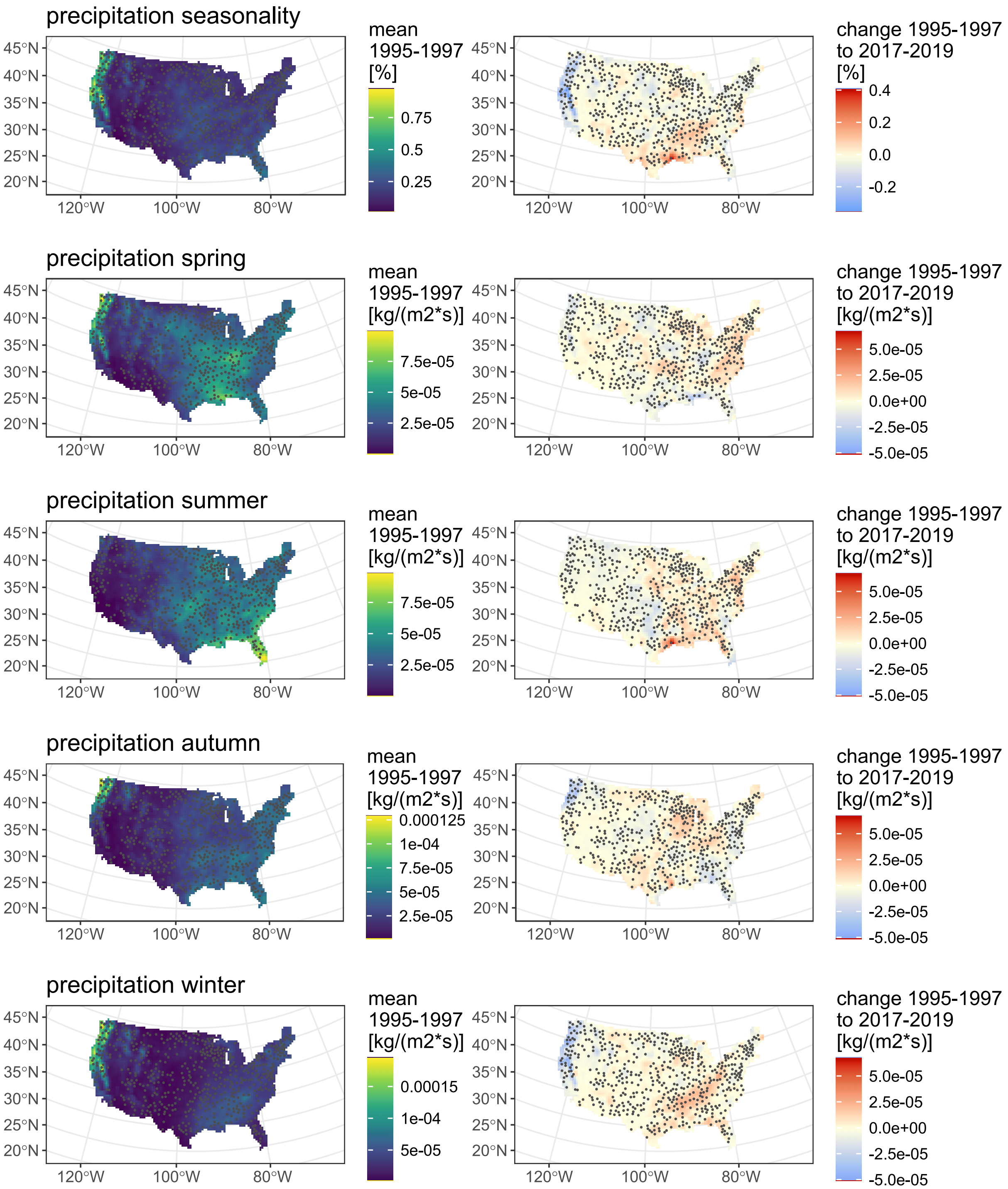


**Figure S 8 continued (2 of 3).**

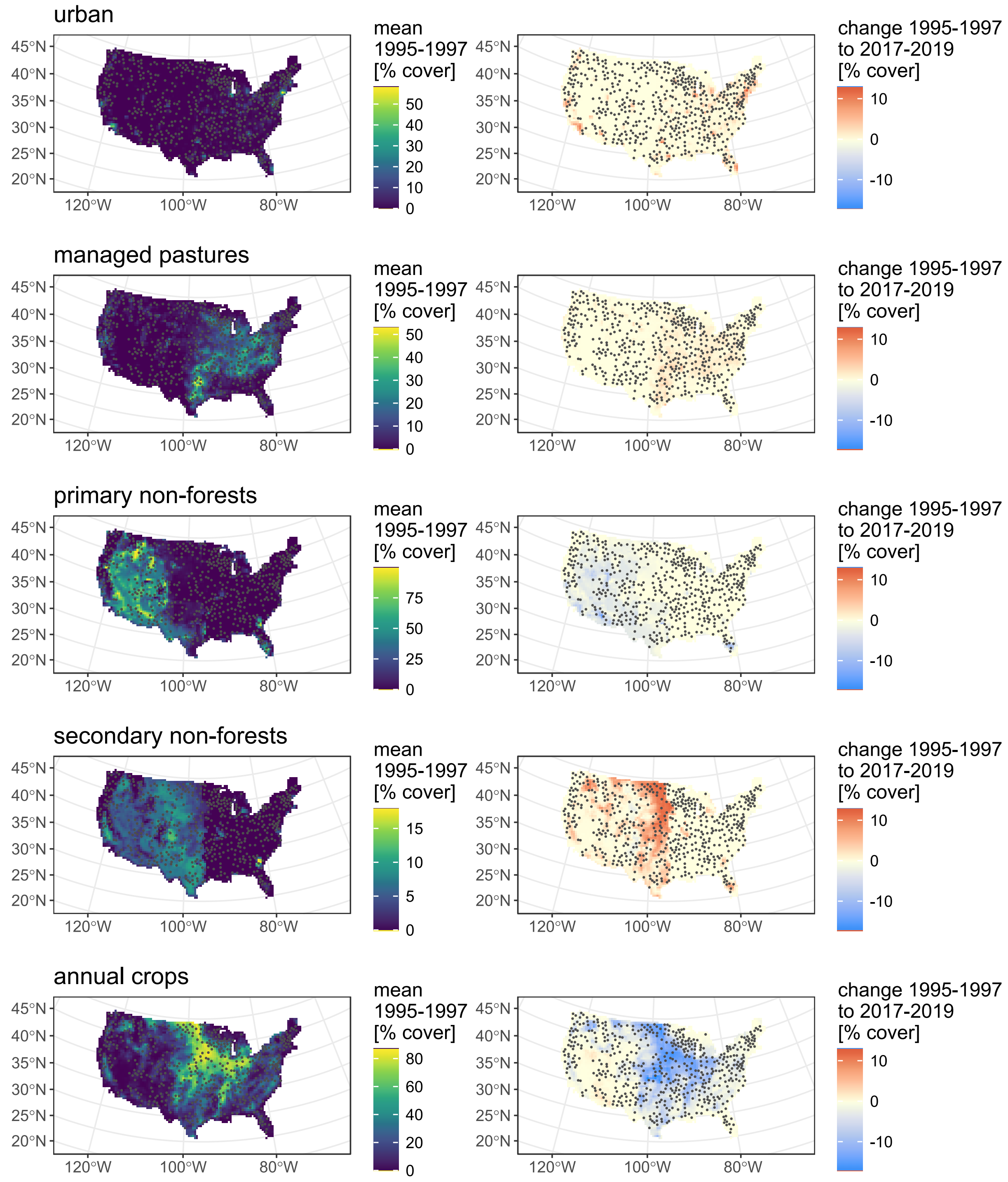


**Figure S 8 continued (3 of 3).**
